# An atlas of anterior *hox* gene expression in the embryonic sea lamprey head: *hox*-code evolution in vertebrates

**DOI:** 10.1101/571448

**Authors:** Hugo J. Parker, Marianne E. Bronner, Robb Krumlauf

**Affiliations:** Stowers Institute for Medical Research, Kansas City, Missouri 64110, USA; Division of Biology and Biological Engineering, California Institute of Technology, Pasadena, California 91125, USA; Department of Anatomy and Cell Biology, Kansas University Medical Center, Kansas City, Kansas 66160, USA

**Keywords:** *Hox* expression, hindbrain segmentation, cranial neural crest, vertebrate evolution, rhombomeres, lamprey, gene regulation, axial patterning

## Abstract

In the hindbrain and the adjacent cranial neural crest (NC) cells of jawed vertebrates (gnathostomes), nested and segmentally-restricted domains of *Hox* gene expression provide a combinatorial *Hox*-code for specifying regional properties during head development. Extant jawless vertebrates, such as the sea lamprey *(Petromyzon marinus),* can provide insights into the evolution and diversification of this *Hox*-code in vertebrates. There is evidence for gnathostome-like spatial patterns of *Hox* expression in lamprey; however, the expression domains of the majority of lamprey *hox* genes from paralogy groups (PG) 1-4 are yet to be characterized, so it is unknown whether they are coupled to hindbrain segments (rhombomeres) and NC. In this study, we systematically describe the spatiotemporal expression of all 14 sea lamprey *hox* genes from PG1-PG4 in the developing hindbrain and pharynx to investigate the extent to which their expression conforms to the archetypal gnathostome hindbrain and pharyngeal *hox-*codes. We find many similarities in *Hox* expression between lamprey and gnathostome species, particularly in rhombomeric domains during hindbrain segmentation and in the cranial neural crest, enabling inference of aspects of *Hox* expression in the ancestral vertebrate embryonic head. These data are consistent with the idea that a *Hox* regulatory network underlying hindbrain segmentation is a pan vertebrate trait. We also reveal differences in hindbrain domains at later stages, as well as expression in the endostyle and in pharyngeal arch (PA) 1 mesoderm. Our analysis suggests that many *Hox* expression domains that are observed in extant gnathostomes were present in ancestral vertebrates but have been partitioned differently across *Hox* clusters in gnathostome and cyclostome lineages after duplication.

## 1. Introduction

*Hox* genes encode a family of highly conserved homeodomain-containing transcription factors that are found in nearly all animal genomes, playing common roles in regulating the specification of positional identities along the anterior-posterior (A-P) axis (Carroll, 1995; Graham et al., 1989). They reside in clusters, with mammals having four paralogous *Hox* clusters, which arose by duplication from a common ancestral complex early in vertebrate evolution (Duboule and Dollé, 1989; Parker and Krumlauf, 2017; Pascual-Anaya et al., 2013). Further duplication and gene loss events have shaped the *Hox* complement across vertebrate lineages (Kuraku and Meyer, 2009; Pascual-Anaya et al., 2013). Based on their sequence similarity and positions within a cluster, vertebrate *Hox* genes are classified into 14 paralogy groups (PG) (Krumlauf, 1994). A regulatory feature of the *Hox* clusters in vertebrates is that during development the timing and domains of *Hox* gene expression along the A-P axis are correlated with their relative gene order along the cluster, a property termed collinearity. Genes within a given *Hox* cluster are all transcribed in the same 5’ to 3’ orientation. *Hox* genes closest to the 3’ end (‘anterior’ *Hox* genes, such as those in PG1) show a tendency to be expressed earlier (temporal colinearity) and more anteriorly (spatial colinearity) than those closer to the 5’ end (Duboule, 2007; Duboule and Dollé, 1989; Kmita and Duboule, 2003). This results in a nested series of *Hox* expression domains, which create combinatorial *‘Hox* codes’ that specify regional properties along the A-P axis in multiple tissues (Mallo et al., 2010).

During embryonic development the vertebrate hindbrain is transiently segmented along the A-P axis into 7 or 8 morphological units, called rhombomeres (r) (Hanneman et al., 1988; Lumsden, 2004; Lumsden and Keynes, 1989). These represent lineage-restricted cellular compartments, which respond to axial patterning signals to create distinct regional identities in each individual segment (Fraser et al., 1990; Marshall et al., 1992). Rhombomeres contain reiterated populations of neurons, which differentiate in a rhombomere-specific manner, resulting in the specialization of morphology, connectivity and function within each segment (Keynes and Lumsden, 1990; Lumsden and Keynes, 1989). The embryonic pharynx also exhibits segmentation, forming an alternating series of pharyngeal arches (PA) and pouches by out-pocketing of the endoderm. Hindbrain segmentation influences craniofacial patterning through cranial neural crest (NC) cells, which delaminate from the neural tube and migrate to the pharyngeal arches in discrete streams (Le Douarin and Kalcheim, 1999). Specific rhombomeres contribute to the different NC streams (Kontges and Lumsden, 1996; Trainor et al., 2002), with signals from surrounding tissues and between rhombomeres influencing NC migratory routes (Golding et al., 2000; Lumsden et al., 1991; Trainor and Krumlauf, 2000a; Trainor and Krumlauf, 2000b; Trainor et al., 2002). Cranial nerves connect each pharyngeal arch to branchiomotor neurons in specific rhombomeres, forming a somatotopic map of the pharyngeal arches in the hindbrain (Lumsden and Keynes, 1989; Oury et al., 2006). Thus, rhombomeres and pharyngeal segments are fundamentally coupled by the migration of cranial NC and by neuronal connectivity between the hindbrain and pharynx.

*Hox* genes are coupled to the gene regulatory network patterning hindbrain segments and NC. A hallmark of *Hox* gene expression in the hindbrain and pharynx is that anterior expression domains correspond tightly with rhombomere and pharyngeal arch boundaries, giving rise to region-specific positional *Hox*-codes in the hindbrain and NC (Hunt et al., 1991; Lumsden and Krumlauf, 1996). Perturbation experiments in jawed vertebrate species have revealed multiple roles for anterior *Hox* genes in hindbrain segmentation, segmental patterning of neurogenesis, and in patterning the skeleton of the head and neck. In the mouse, *Hoxa1* is required early in hindbrain development for the formation of r5 (Chisaka et al., 1992; Dollé et al., 1993; Mark et al., 1993), while *Hoxb1* influences neurogenesis in r4 (Goddard et al., 1996; Studer et al., 1996). In mice and zebrafish that lack *Hoxb1,* r4 neurons adopt the characteristics of those in r2, exhibiting altered migration and pathfinding of motoneurons (McClintock et al., 2002; Studer et al., 1996). In an analogous manner, *Hox* genes also have complex inputs into NC. Loss of *Hoxa1* and *Hoxb1* in the mouse neural tube results in a failure to form the r4-derived NC which migrates into PA2 (Gavalas et al., 1998; Gavalas et al., 2001). In diverse vertebrate models, loss of *Hoxa2* leads to a partial transformation of PA2 skeletal derivatives into PA1-like structures (Baltzinger et al., 2005; Gendron-Maguire et al., 1993; Hunter and Prince, 2002; Rijli et al., 1993), while ectopic expression of *Hoxa2* in PA1 leads to duplication of PA2 derivatives (Grammatopoulos et al., 2000; Kitazawa et al., 2015; Pasqualetti et al., 2000). Thus, *Hoxa2* acts as a selector gene for specifying PA2 derivatives, while Hox paralogy group (PG) 1 genes regulate steps in the formation of NC.

*Hox* segmental patterning roles in the hindbrain and NC appear to be widely conserved across jawed vertebrates, based on functional studies in multiple species (Baltzinger et al., 2005; Grammatopoulos et al., 2000; Hunter and Prince, 2002; McClintock et al., 2002). Expression studies in dogfish, a cartilaginous fish, root their ancestry at least to the base of the jawed vertebrates (Oulion et al., 2011). This deep ancestry is also reflected by the sequence conservation of *Hox* enhancers that modulate segmental expression and by the conservation of *Hox-*responsive enhancer elements associated with downstream target genes (Kim et al., 2000; McEwen et al., 2009; Parker and Krumlauf, 2017; Parker et al., 2011; Parker et al., 2014b; Ravi et al., 2009). Invertebrate chordates, such as amphioxus (a cephalochordate) and *dona* (a urochordate), display nested and co-linear *Hox* expression along the A-P neuraxis. The conservation of vertebrate-like retinoic acid response elements in the amphioxus *Hox* cluster suggests that ancestral chordates used in part an RA-Hox regulatory circuitry to generate nested A-P *Hox* expression in neural patterning (Manzanares et al., 2000; Wada et al., 2006). However, unlike vertebrates, invertebrate chordates lack rhombomeric segmentation and definitive NC. This raises two evolutionary questions: First, when in vertebrate evolution did these segmental *Hox* roles evolve? Second, how have these roles diverged between vertebrate lineages? Lamprey and hagfish belong to a lineage of jawless extant vertebrates (cyclostomes), which diverged early in vertebrate evolution from the lineage leading to the jawed vertebrates (gnathostomes), making them important species for addressing questions about early vertebrate evolution and diversification (Shimeld and Donoghue, 2012).

The ancestor of extant vertebrates is inferred, based on parsimony, to have had 4 *Hox* clusters, arising from a single ancestral chordate cluster through genomic duplication events in early vertebrates. Genomic analyses in two lamprey species - sea lamprey *(Petromyzon marinus)* and the closely related Arctic lamprey (*Lethenteron camtschaticum) -* revealed each to possess 6 *hox* clusters, indicative of additional duplication event/s in the lamprey/cyclostome lineage (Figure1) (Mehta et al., 2013; Smith et al., 2018). This raises the prospect that roles for *hox* genes could have diversified in lamprey after these *hox* cluster duplications, with duplicated *hox* genes potentially being associated with anatomical novelties. To date, detailed expression analyses have been reported for only 3 anterior (PG1-4) *hox* genes in sea lamprey (Parker et al., 2014a), and for 5 such genes in Arctic lamprey (Takio et al., 2007). Sea lamprey was found to have transient rhombomere-restricted *hox* expression in the hindbrain and nested *hox* domains in the NC, similar to gnathostomes (Parker et al., 2014a; Takio et al., 2004). However, given that the sea lamprey has 14 anterior *hox* genes, the expression domains of the majority of lamprey *hox* PG1-4 genes are yet to be characterized, so it is unknown whether they are coupled to hindbrain segmentation and NC. Thus, the extent to which *hox* expression in the head is conserved or divergent between jawed and jawless vertebrates is still unclear, calling for a more comprehensive analysis of lamprey *hox* gene expression.

In this study, we systematically describe the spatiotemporal expression of all 14 lamprey anterior *hox* genes in PG1-4 in the developing hindbrain and pharynx. We address the extent to which their expression conforms to the archetypal gnathostome hindbrain and pharyngeal *hox-*codes. In the context of lamprey/cyclostome-specific *hox* cluster duplications, we investigate whether the resulting paralogues exhibit equivalent or divergent patterns of expression. Finally, these expression patterns are used as a basis to infer shared and divergent aspects of *hox* cranial patterning between jawed and jawless vertebrates.

## 2. Materials and Methods

### 2.1 Lamprey embryos

Lamprey husbandry and embryo collection was performed as previously described (Nikitina et al., 2009; Parker et al., 2014a), with embryos being staged according to Tahara (Tahara, 1988), fixed in MEMFA, and dehydrated in 100% ethanol for storage at −20°C. This study was conducted in accordance with the Guide for the Care and Use of Laboratory Animals of the National Institutes of Health and protocols were approved by the Institutional Animal Care and Use Committee of the California Institute of Technology (lamprey, Protocol #1436-17).

### 2.2 Cloning of cDNA for in situ hybridization probes

*In-situ* probes were designed based on predicted gene sequences in the sea lamprey germline genome assembly (gPMAR100)(Smith et al., 2018), with care taken to avoid repetitive elements. Probe sequences were amplified from *P. marinus* genomic DNA or from st18-26 embryonic cDNA by PCR using KOD Hot Start Master Mix (Novagen). 3’ rapid amplification of cDNA ends (RACE) was performed for *wnt1* using the GeneRacer Kit (Thermo Fisher Scientific). PCR products were cloned into the *pCR4-TOPO* vector (Thermo Fisher Scientific) and sequenced. The following PCR primers were used for amplifying probe templates, with probe lengths given:

*wnt1* (729bp, partial exon and 3’UTR) F: 5’-GAACTGCACGCGGGTGGAGACTGT-3’; R: GeneRacer 3’ Nested Primer.

*hoxβ1* (674bp, 3’UTR fragment) F: 5’-ATGCTCCCTCAACTCCATCC-3’; R: 5’-TGACCTCTTCTCGCATGTAAGA-3’.

*hoxε1* (338bp, partial exon 2) F: 5’-GCTGCTTCCACCAACAGG-3’; R: 5’-GAACCCCTTCGCCGAGAC-3’.

*hoxζ1* (556bp, 3’UTR fragment) F: 5’-AGACATCCGGGCAATCGATT-3’; R: 5’-ATCGCTACTTCGCCAAATCG-3’.

*hoxδ2* (585bp, partial exon 2) F: 5’-ACCTCTGCGCGACTCCTC-3’; R: 5’-CCAGACCTCCTCCTCCTCT-3’.

*hoxδ3* (359bp, partial exon 2) F: 5’-GAGAACTCGTGCGGTGG-3’; R: 5’-TTGCCCAAACCGTGCAG-3’.

*hoxζ*3 (321bp, partial exon 2) F: 5’-TACCACCTCGTCGTCCAC-3’; R: 5’-GACAGCCTCGACCCCAAA-3’.

*hox*α*4* (301bp, partial exon 1-2) F: 5’-CTGAAGCAGCCGGTCGTG-3’; R: 5’-TGGACGAGGCTGTGTTCAAT-3’.

*hoxβ4* (403bp, partial exon 1-2) F: 5’-AGCAGCAGGGACACTTGAT-3’; R: 5’-GAACGGATCTTGGTGTTGGG-3’.

*hoxγ4* (267bp, partial exon 1-2) F: 5’-ACCCGTGGATGAAGAAGGTA-3’; R: 5’-TCACCTTGGTGTTCGGTAGT-3’.

*hoxδ4* (382bp, partial exon 2) F: 5’-CCAGGGACACGAGACCAAA-3’; R: 5’-GCTGGGCCTAACTCCTCAAA-3’.

*hoxε4* (338bp, partial exon 2) F: 5’-CAACTATATCGGCGGGGAGT-3’; R: 5’-TGCTACTACCATTGCTGCTG-3’.

*hoxζ4* (382bp, partial exon 1-2) F: 5’-GCGGTGACTTCAACCATCAA-3’; R: 5’-GCAGCTTGTGGTCCTTCTTC-3’.

*krox20, hoxα2, hoxα3* probe sequences are as previously reported(Parker et al., 2014a).

### 2.3 In situ hybridization

Digoxygenin-labelled probes were generated by standard methods and purified using the MEGAclear Transcription Clean-up Kit (Ambion). Lamprey wholemount *in situ* hybridization was performed as described previously (Nikitina et al., 2009; Sauka-Spengler et al., 2007), with the following amendments to the protocol: methanol-stored embryos were first transferred into ethanol and left overnight prior to rehydration; a treatment of 0.5% acetic anhydride in 0.1M triethanolamine was added after proteinase K digestion. Hybridization was performed at 70°C for each probe. Embryos were cleared either by using a glycerol series followed by imaging in 100% glycerol, or by using a 1:2 ratio of benzyl alcohokbenzyl benzoate followed by mounting in Permount (Fisher Scientific) on microscope slides for imaging.

### 2.4 Sectioning

After *in situ* hybridization, selected embryos were transferred to 30% sucrose in phosphate-buffered saline, embedded in O.C.T compound (VWR), and cryo-sectioned to 10μm-thick sections.

### 2.5 Imaging

Images of BABB-cleared embryos were taken using a Zeiss Axiovert 200 microscope with an AxioCam HRc camera and AxioVision Rel 4.8.2 software. Glycerol-cleared embryo images were taken using a Leica MZ APO microscope with a Lumenera Infinity 3 camera and Infinity Analyze software. Sections were imaged using a Zeiss Axiovert 200 microscope with a Lumenera Infinity 3 camera and Micro-Manager 1.4.22 software. Images were cropped and altered for brightness and contrast using Adobe Photoshop CS5.1.

### 2.6 Data -Availability

Original data underlying this manuscript can be accessed from the Stowers Original Data Repository at [http://odr.stowers.org/websimr/]

## 3. Results

### 3.1 The lamprey hox complement

The sea lamprey and the Arctic lamprey each have 42 *hox* genes arranged in 6 clusters and 14 paralogy groups, compared to mouse with 39 *hox* genes across 4 clusters and 13 PG (Fig. 1) (Mehta et al., 2013; Smith et al., 2018). Within PG1-4, the *Hox* gene content is very similar between lamprey and mouse: both have 3 PG1, 2 PG2 and 3 PG3 genes, while lamprey has 6 PG4 genes compared to 4 in mouse (Fig. 1). Phylogenetic analyses could not resolve direct orthology between specific lamprey and gnathostome *hox* clusters (Mehta et al., 2013; Smith et al., 2018). Synteny analysis based on the retention of paralogous genes between lamprey *hox*-bearing chromosomes found significant similarity in gene content between chromosomes containing the lamprey −β and −ε clusters, and between those containing the −α and −δ clusters (Smith et al., 2018). This suggests that these pairs of chromosomes arose from duplication event/s that occurred in the cyclostome/lamprey lineage, after the split from the lineage leading to gnathostomes (Fig. 1). It has been suggested, based on parsimony, that the ancestor of all extant vertebrates had 4 *Hox* clusters, resulting from duplication events in an early vertebrate lineage, consistent with a recent reconstruction of vertebrate chromosomal evolution (Smith et al., 2018). Taken together, this leads to a scenario in which the common ancestor of gnathostomes and cyclostomes had 4 *Hox* clusters, with additional chromosome-scale (or possibly whole-genome) duplications occurring in the cyclostome/lamprey lineage, resulting in the 6 *Hox* clusters of extant lampreys. Of the anterior *hox* genes (PG1-4) in sea lamprey, only 3 have had their expression characterized by *in-situ* hybridization (Fig. 1 - lilac shading).

**Figure 1:**
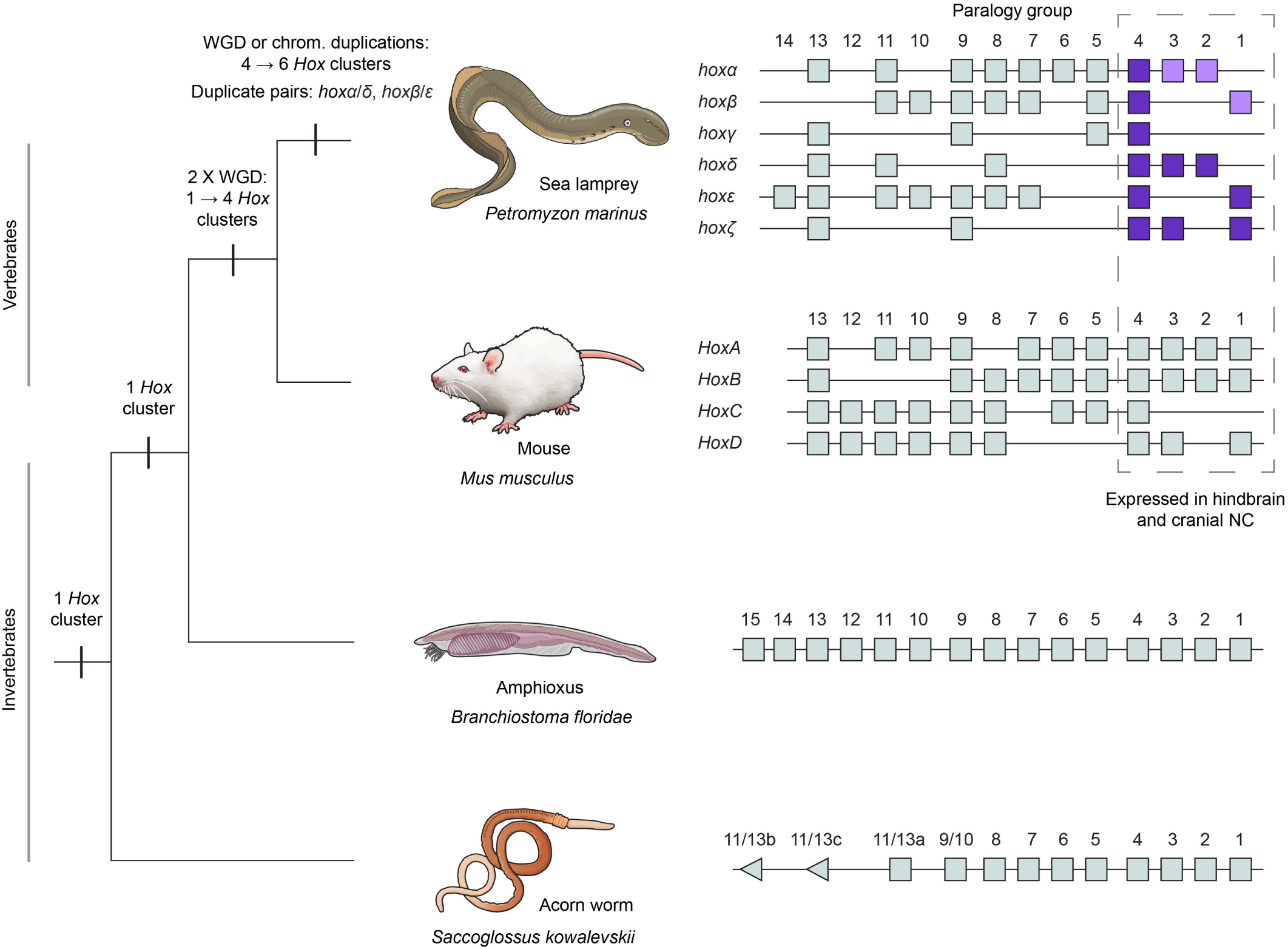
*Hox* clusters of selected deuterostomes. A phylogeny of selected deuterostomes, showing the characterised *Hox* clusters for given species. The duplication events that are inferred to have shaped the *Hox* complement of these species are indicated. These include whole genome duplication/s (WGD) in the early vertebrate lineage and WGD or chromosomal duplications in the cyclostome lineage leading to lamprey. The Hox clusters are depicted with the direction of transcription from left to right. Acorn worm hox11/13b and 11/13c show opposite direction of transcription to the rest of the *hox* cluster. For sea lamprey, *hox* genes previously characterised by *in-situ* hybridisation are shaded in lilac, and those characterised for the first time in this study shaded in purple.

### 3.2 The segmental plan of the lamprey embryonic hindbrain and pharynx

At st23.5, at least six rhombomeres can be demarcated by gene expression in the lamprey hindbrain, with *wnt1* expressed in the midbrain and abutting the midbrain-hindbrain boundary, *krox20* in r3/r5, *hoxζ4* (a PG4 gene, described in more detail below) posterior to and abutting the r6/r7 boundary, and the anterior border of *hoxα2* marking the r1/r2 boundary (Fig. 2A-D). *hoxβ1* and *hoxα3* exhibit discrete stripes of rhombomere-restricted expression at this stage, in r4 and r5 respectively, as previously shown (Fig. 2E-H) (Parker et al., 2014a).

**Figure 2:**
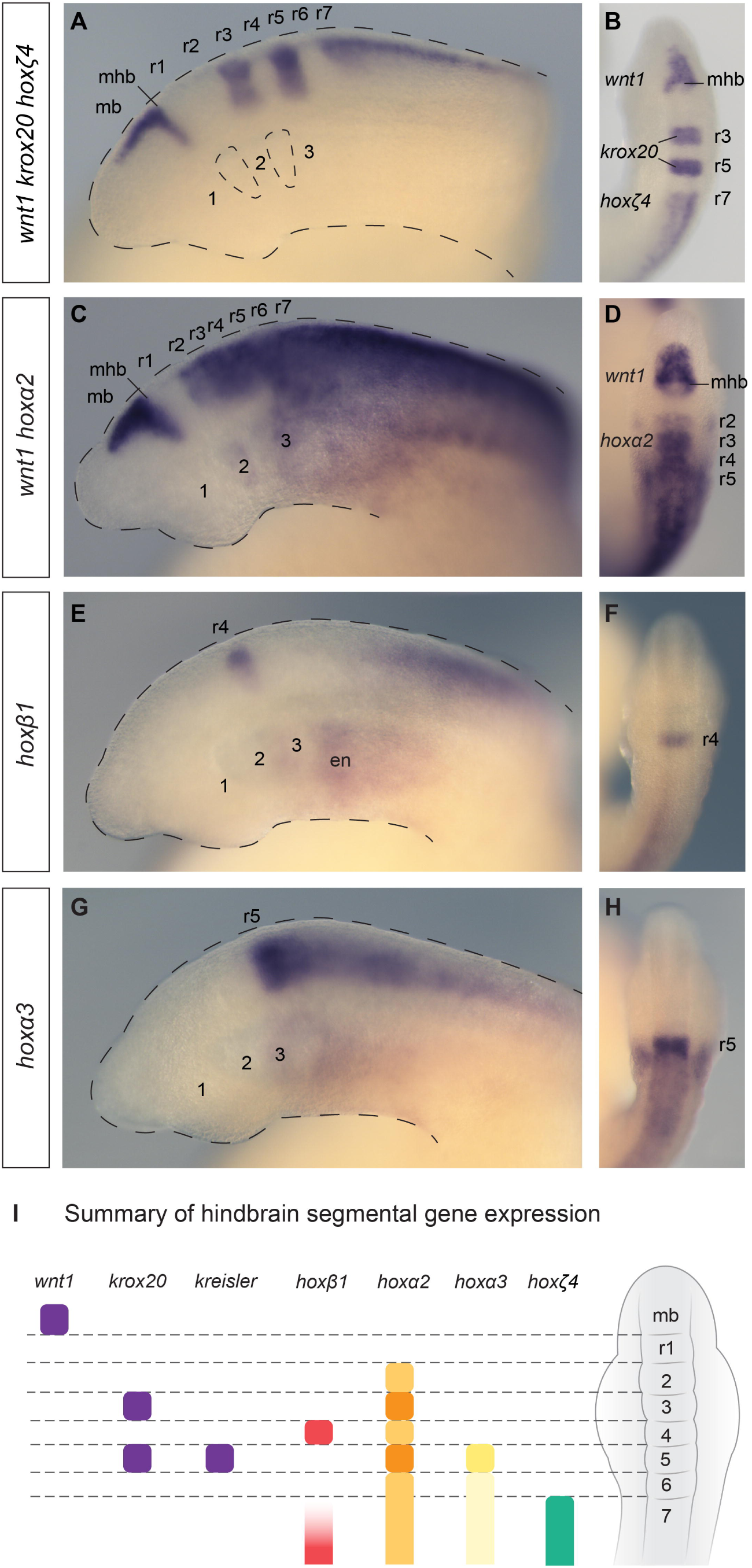
The lamprey hindbrain segmental plan and segmental hox expression. Lateral (A,C,E,G) and dorsal (B,D,F,H) views of st23.5 lamprey embryos are shown. (A-B) A triple *in-situ* hybridisation against *wnt1, krox20* and *hoxζ4* (all purple) demarcates hindbrain segments in the neural tube, *wnt1* marks the caudal limit of the midbrain (mb), revealing the midbrain-hindbrain boundary (mhb), while *krox20* marks r3 and r5, and *hoxζ4* is expressed posterior to the r6/r7 boundary. Rhombomeres (r1-r7) and pharyngeal arches (1-3) are annotated, and the head and pharyngeal pouches are outlined. (C-D) A double in-situ hybridisation against *wnt1* and *hoxα2,* showing segmental *hoxα2* expression in the hindbrain posterior to the r1/r2 boundary and *wnt1* in the midbrain. *hoxα2* is also expressed in the developing pharyngeal arches, posterior to PA1. (E-F) *hoxβ1* is expressed in r4 and in the posterior hindbrain/spinal cord, as well as in the pharyngeal endoderm (en). (G-H), *hoxα3* shows an elevated stripe of expression in r5, with lower expression levels in the neural tube posterior to r5. Expression is also seen in the pharyngeal arches, posterior to PA2. (I) A depiction of a dorsal view of a st23.5 lamprey embryo, summarising the segmental gene expression domains in the neural tube shown in (A-H), which together demarcate r1-r7. *kreisler* expression was previously characterized in r5 (Parker et al., 2014a).

Lamprey pharyngeal segmentation occurs between st21-26, as the pharynx is progressively segmented into a series of pharyngeal arches and pouches, ultimately comprising 8 pharyngeal arches by st26. From st23, *hoxα2* is visible in the pharyngeal arches, with an anterior limit in PA2 (Figure2C), while *hoxα3* has an anterior limit in PA3 (Fig. 2G). Together, these segmental patterns in the hindbrain and pharynx provide a topographical and temporal framework in which to analyze the expression of the anterior *hox* genes during lamprey head development (Fig. 2I).

### 3.3 hox PG1 expression

We first investigated the expression of the three lamprey PG1 genes – *hoxβ1, hoxε1* and *hoxζ1.*In gnathostomes, PG1 genes are the earliest *Hox* genes to be expressed in the neuroepithelium, so we investigated their expression during early lamprey development. We detected differential timing of onset in the neuroepithelium between these genes, with *hoxζ1* and *hoxε1* first detectable at st17 in broad and overlapping domains that develop clear anterior boundaries by st18 (Fig. 3A). These domains persist through st20, with *hoxβ1* expression in the neural plate emerging by st19. At st20, all three PG1 genes show similar anterior borders with high levels of neural expression in putative r4. *hoxβ1* resolves into a distinct anterior stripe, while *hoxε1* and *hoxζ1* retain lower levels of expression posterior to r4. Expression adjacent to the neural plate is also seen for *hoxβ1* and *hoxε1.* At later stages, st21-26, *hoxβ1* remains as a restricted stripe in r4, with additional expression in the posterior hindbrain and spinal cord, as previously reported (Fig. 3B) (Parker et al., 2014a). In contrast, *hoxε1* and *hoxζ1* expression is lost from r4, but persists more posteriorly in the neural tube, with *hoxζ1* expression then disappearing from the neural tube by st25 (Fig. 3B). *hoxβ1* and *hoxε1* also show expression in bilateral clusters of cells in the region of the forebrain/midbrain boundary at st24-25, which may be homologous to that characterised for gnathostome PG1 genes in a similar domain (McClintock et al., 2003).

**Figure 3:**
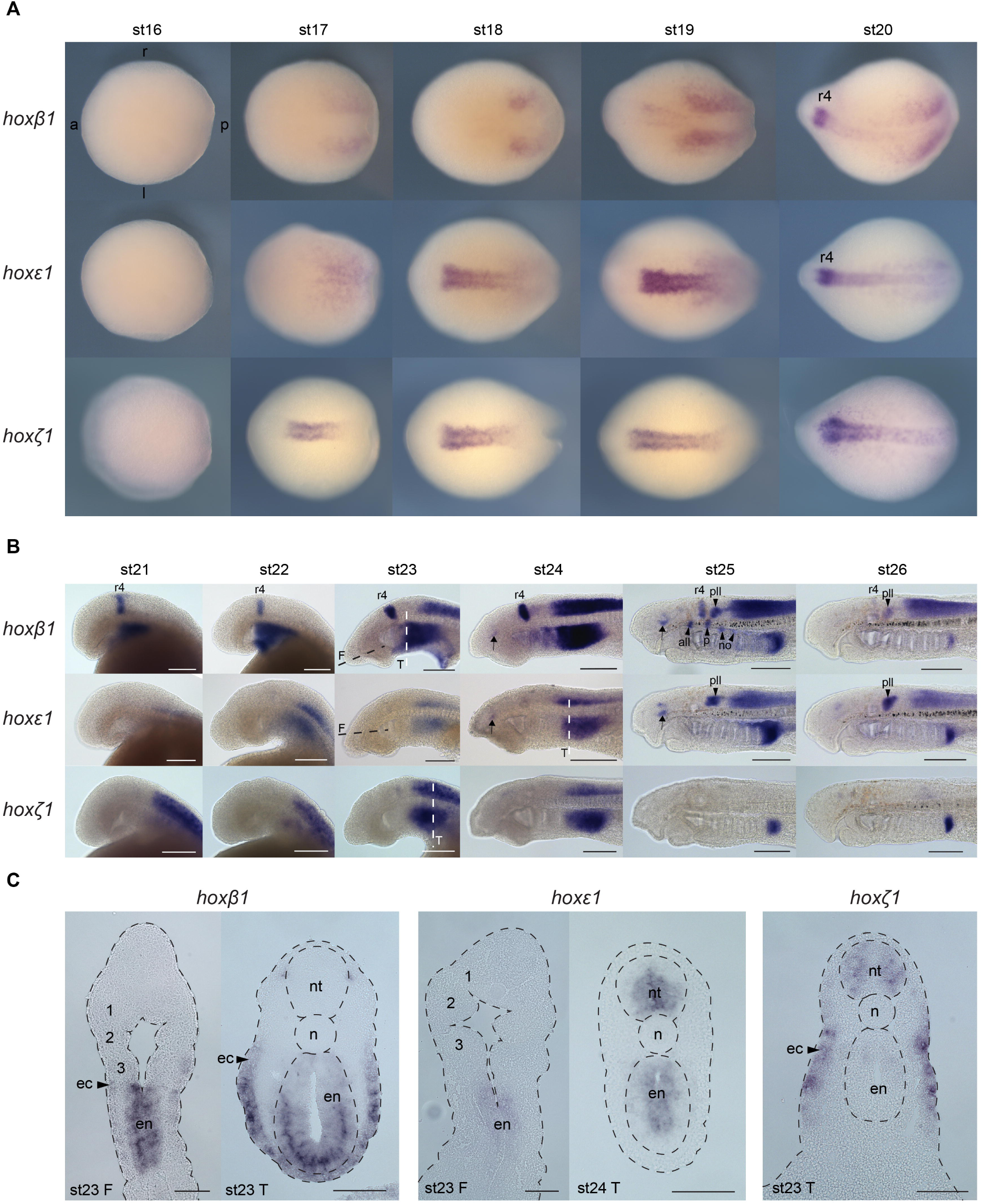
Expression of lamprey *hox*PG1 genes in the developing head. (A) Dorsal views of st16-st20 embryos with expression of *hox*PG1 genes revealed by *in situ* hybridisation. The anterior (a), posterior (p), left (l) and right (r) sides are annotated in the top-left image. (B) Expression of *hox*PG1 genes in st21-26 embryos, shown in lateral view with anterior to the left. Arrows mark neurons in the forebrain/midbrain. Arrowheads label cranial ganglia: all, anterior lateral line ganglion; no, nodose ganglion; p, petrosal ganglion; pll, posterior lateral line ganglion. (C) Frontal (F) and transverse (T) sections of st23-24 embryos after in situ hybridisation. The approximate planes of section are indicated in the lateral views shown in panel B. The developing pharyngeal arches are annotated (1-3) on the frontal sections. Scale bars: 200μm (B); 100μm (C). ec, ectoderm; en, endoderm; n, notochord; nt, neural tube; r, rhombomere.

In the pharynx, *hoxβ1* is prominently expressed in the endoderm and ventral ectoderm from st21, (Fig. 3B,C). Expression is temporally dynamic in both tissues, regressing posteriorly during pharyngeal segmentation to remain positioned posterior to the most recently formed pharyngeal pouch. *hoxε1* also displays similar endodermal expression in the pharynx, with *hoxζ1* expressed in the posterior pharyngeal ectoderm (Fig. 3B,C). *hoxβ1* and *hoxε1* are expressed in the cranial ganglia from st25 - both genes in the posterior lateral line ganglion, *hoxβ1* in the anterior lateral line, petrosal and nodose ganglia (Fig. 3B).

### 3.4 hox PG2 expression

In the neural tube, *hoxα2* is expressed in presumptive r3 and r5 from st21 and has lower levels of expression in r4 and posterior to r5 at that stage (Fig. 4). By st22, expression is also seen in r2 such that prominent rhombomeric stripes are visible in r2-r5. From st24 onwards, expression in the hindbrain and spinal cord persists, with an anterior limit in r2, but the rhombomere-restricted stripes of expression become less clear. *hoxδ2* expression is detected in restricted domains within presumptive r5 and in dorsal r3 from st21, which persists across our developmental time-course (Fig. 4). Additional expression of lower intensity is also seen in the neural tube posterior to r5, with a dorsally-restricted domain caudal to r5 visible at st25-26.

**Figure 4:**
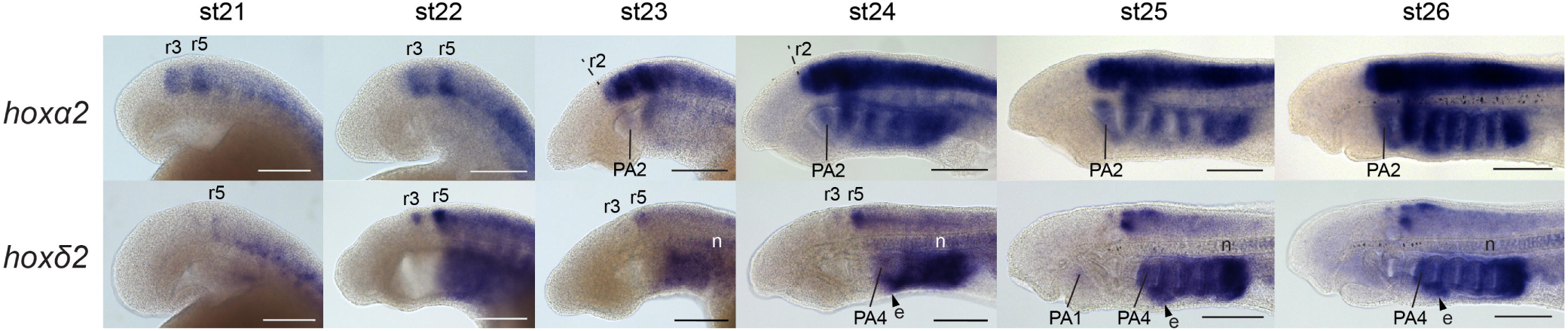
Expression of lamprey *hoxPGl* genes in the developing head. Expression of *hox*PG2 genes in st21-26 embryos, shown in lateral view with anterior to the left. Scale bars: 200μm. e, endostyle; n, notochord; PA, pharyngeal arch; r, rhombomere.

In the pharynx, *hoxα2* is expressed in the pharyngeal arches from st23 and is maintained through later stages, with an anterior limit in PA2. At st25, this expression is prominent in the NC-derived mesenchyme, as well as in the pharyngeal arch mesoderm, as revealed by frontal sectioning (see Fig. 7B). *hoxδ2* expression in the pharynx is seen from st22 and persists to later stages, with an anterior limit at st24 in the caudal half of the third pharyngeal pouch. We also observed transient, faint signal in the first pharyngeal arch from st25-26. Frontal sections at st26 show that this PA1 expression is mesodermal (see Fig. 7B), while the caudal pharyngeal expression is in the pharyngeal endoderm (pharyngeal pouch 3 to posterior) and in the mesenchyme of PA6-8. Expression was also detected for *hoxδ2* in the caudal extent of the developing endostyle, posterior to PA4 at st24-26, as well as in the notochord from st23-26 (Fig. 4).

### 3.5 hox PG3 expression

The PG3 genes show nested expression in the developing hindbrain, with offset anterior boundaries (Fig. 5). *hoxα3* is expressed at a high level in r5 at st22, with lower expression detected in the neural tube posterior to r5. By st23, additional weak expression is detectable in r4. At these stages, *hoxδ3* is expressed posterior to the r5/r6 boundary, and *hoxζ3* posterior to the r6/r7 boundary, as revealed by comparison with *krox20* in r3/r5 (see Fig. 7A). These patterns are temporally dynamic - from st24 onwards they break from rhombomeric registration, with each gene showing anterior expression boundaries that are non-uniform along the dorso-ventral axis. For example, at st25-st26, *hoxδ3* signal is visible in the hindbrain with a sharp anterior border that aligns with the anterior side of PA4, except for a small domain in the dorsal hindbrain that protrudes rostrally from this border. Pharyngeal expression is detected for *hoxα3* and *hoxδ3* but not for *hoxζ3* (Fig. 5). This is visible from st23, resolving into nested domains in the pharyngeal arch mesenchyme by st26: *hoxα3* in PA3-8 and *hoxδ3* in PA4-8 (see Fig. 7B).

**Figure 5:**
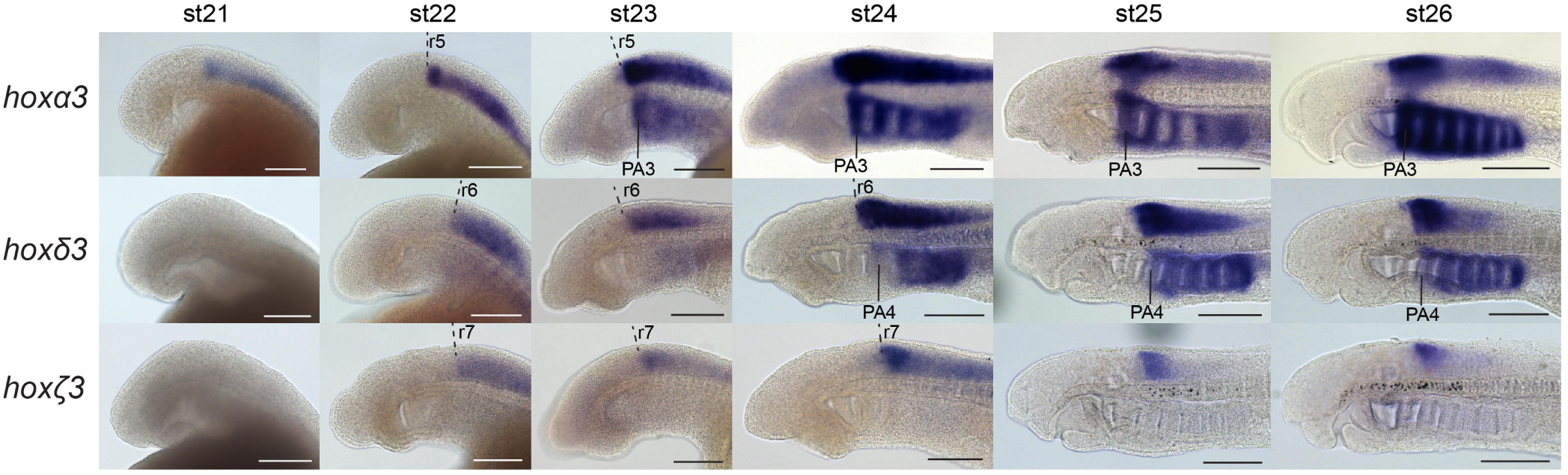
Expression of lamprey *hoxPG3* genes in the developing head. Expression of *hox*PG3 genes in st21-26 embryos, shown in lateral view with anterior to the left. Scale bars: 200μm. PA, pharyngeal arch; r, rhombomere.

### 3.6 hox PG4 expression

We detected expression of each of the 6 PG4 genes in the neural tube across our developmental time-course (Fig. 6). We compared their anterior borders of expression using *krox20* expression, to mark r3 and r5 (Fig. 7A), and by using the pharyngeal arches as landmarks (Fig. 6). This analysis reveals that these genes vary in their precise spatial domains and temporal dynamics. At st24, *hoxα4* and *hoxζ4* have clear anterior expression borders at the same position, aligning with the anterior face of PA4. Comparison with *krox20* expression shows that this domain is caudal to r5 by approximately one rhombomere-length and is thus likely to represent the r6/r7 boundary (Fig. 7A). The other PG4 genes have anterior expression borders that are posterior to this region in the caudal hindbrain. The anterior expression limits in the neural tube are temporally dynamic for some of these PG4 genes: *hoxα4* expression aligns anteriorly with PA4 at st24 but with PA6 at st26, while that of *hoxβ4* aligns with PA4 at st24 and with PA5 at st26 (Fig. 6). In other cases, such as *hoxζ4,* the anterior boundary is maintained across this time-course, appearing to retain a tight rhombomeric registration. *hoxγ4, hoxδ4,* and *hoxε4* each show expression profiles that change along the dorsal-ventral axis across this time-course, with expression in the dorsal neural tube spreading more rostrally in each case, perhaps encompassing specific neuronal populations.

**Figure 6:**
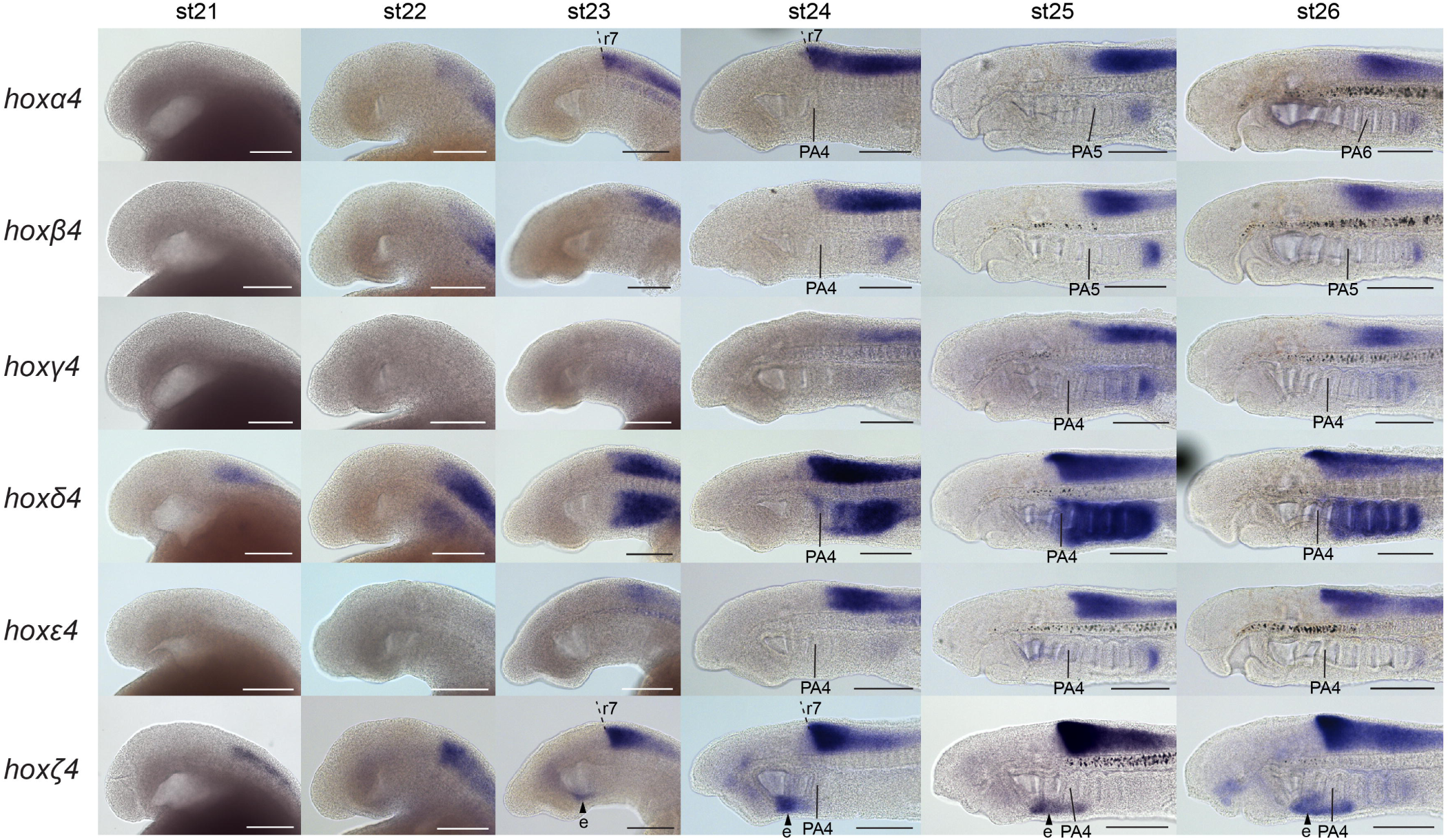
Expression of lamprey *hox*PG4 genes in the developing head. Expression of *hox*PG4 genes in st21-26 embryos, shown in lateral view with anterior to the left. Neural expression is seen in the posterior hindbrain and/or spinal cord for each gene. To facilitate comparison of this neural expression across time and between genes, the pharyngeal arches (PA) that are adjacent to the anterior neural expression boundaries are labelled. Scale bars: 200μm. e, endostyle; PA, pharyngeal arch; r, rhombomere.

**Figure 7:**
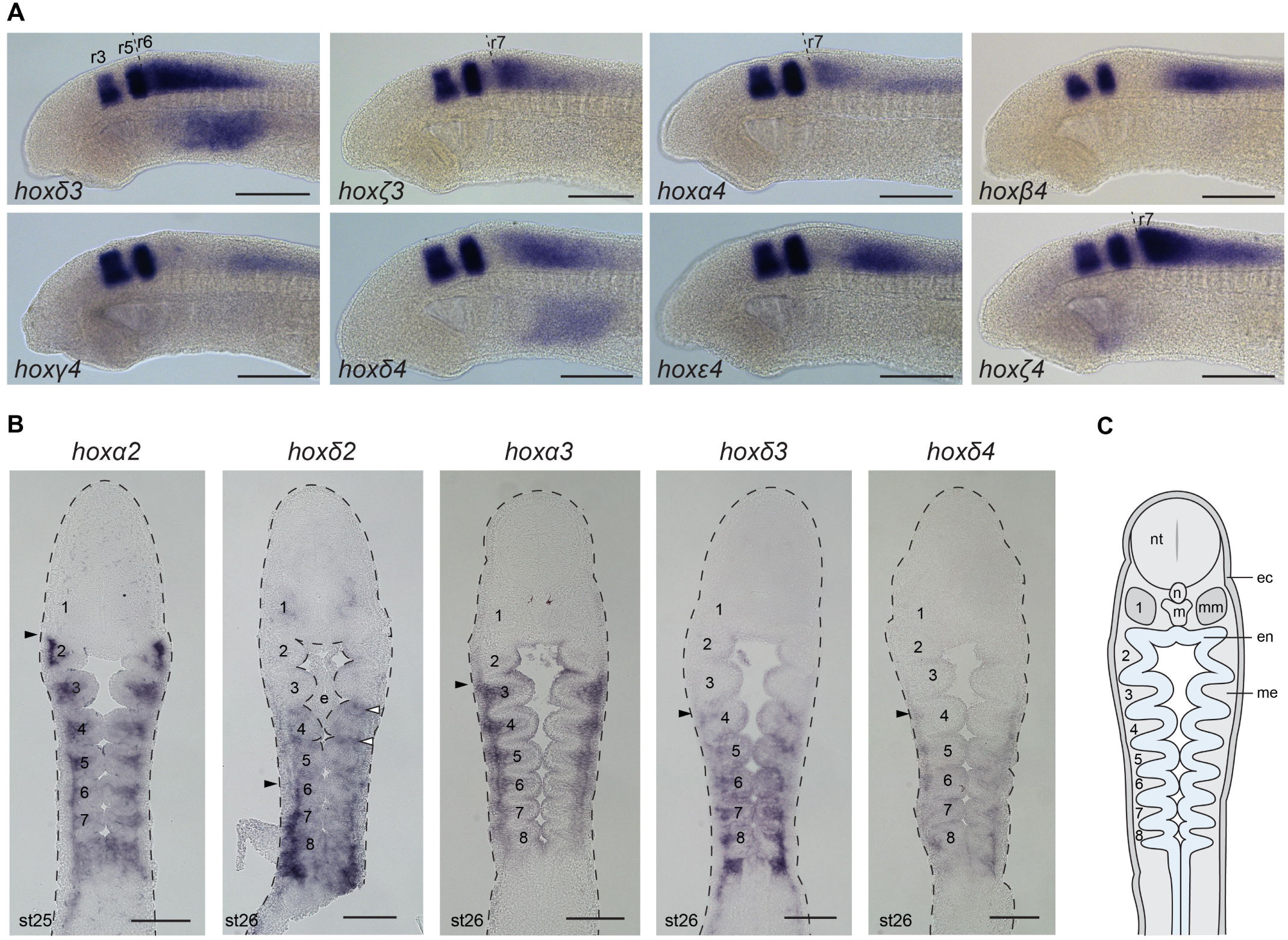
Rhombomeric *hoxPG3-4* expression and pharyngeal expression of selected lamprey *hoxPG1-4* genes. (A) Double in situ hybridization of *hox* genes from PG3-4 with *krox20,* to resolve rhombomeric expression domains. *krox20* expression is in r3 and r5. For the PG3-4 genes with clear rhombomeric boundaries, they are indicated (dashed line). *hoxα3* expression in r5 was previously characterized and so is not shown (Parker et al., 2014a). (B) Frontal sections at st25-26, revealing pharyngeal hoxPG2-4 expression domains. Black arrowheads indicate anterior expression limits in the neural crest-derived pharyngeal arch mesenchyme. White arrowheads mark *hoxδ2* expression in pharyngeal pouch endoderm. Pharyngeal arches are numbered (1-8). (C) A schematic frontal section of a st26 lamprey embryo indicating the different tissue layers, ec, ectoderm; en, endoderm; m, mouth; me, mesenchyme; mm, mandibular mesoderm; n, notochord; nt, neural tube.

Expression is visible for each of the *hox*PG4 genes within the pharynx (Figure6). For *hoxα4, hoxβ4,* and *hoxε4,* this signal was only detectable in the most caudal extent of the pharynx. *hoxγ4* shows faint signal in a gradient of increasing intensity from PA5 caudally, while *hoxδ4* expression is visible in the mesenchyme of PA4-8 (Figure7B). *hoxζ4* was detectable in the developing endostyle from st23 onwards, but expression in other pharyngeal domains was not seen for this gene.

## 4. Discussion

We have characterised the expression of the 14 *Hox* PG1-4 genes in the developing lamprey head to address two primary questions: when did segmental *Hox* domains evolve in vertebrate evolution and how have they diversified between vertebrate lineages? We find many similarities in *Hox* expression between lamprey and gnathostome species, particularly in rhombomeric domains during hindbrain segmentation and in the cranial neural crest, enabling inference of aspects of *Hox* expression in the ancestral vertebrate embryonic head. We also observe differences, including variation in hindbrain domains at later stages, as well as expression in the endostyle and in PA1 mesoderm. Considering the *Hox* cluster duplications that preceded the cyclostome-gnathostome divergence, comparison of *Hox* expression and cluster organization between lamprey and gnathostomes suggests that ancestral vertebrate *Hox* functions have been largely retained in lamprey and gnathostomes but have been partitioned differently across duplicated clusters in each lineage. This is consistent with the observation that a *Hox* regulatory network underlying hindbrain segmentation is conserved to the base of vertebrates (Parker et al., 2014a).

### 4.1 The hox repertoire of lamprey and its relationship to gnathostome Hox clusters

The two lamprey species examined to date both have 6 *Hox* clusters and appear to share an identical *Hox* gene complement, reflecting their close phylogenetic relationship (Kuraku and Kuratani, 2006; Mehta et al., 2013; Pascual-Anaya et al., 2018; Smith et al., 2018). Despite lamprey having 6 *Hox* clusters compared to 4 in most tetrapods, paralogue loss has resulted in the total number of *Hox* genes being similar between these taxa: 43 in lamprey and 39 in mouse (Fig. 1). For PG1-4, lamprey and mouse have both retained a remarkably similar number of genes in each paralogy group. It remains unclear precisely how the 6 lamprey *Hox* clusters relate to the 4 *Hox* clusters that were presumably present in the common ancestor of gnathostomes, and to the 4 clusters in mouse, since phylogenetic analyses could not resolve 1:1 orthology between lamprey and gnathostome *Hox* genes/proteins (Mehta et al., 2013; Pascual-Anaya et al., 2018; Smith et al., 2018). This does not necessarily imply that lamprey and gnathostome *Hox* clusters arose from independent duplication events, since ancient duplication, when followed quickly by lineage separation and subsequent divergence, coupled with species-specific patterns of codon and amino acid usage, could weaken the signal of these phylogenetic events (Qiu et al., 2011). Indeed, recent reconstructions based on comparisons of gene order at the chromosomal level between vertebrate species are consistent with a model in which the ancestor of cyclostomes and gnathostomes had 4 *Hox* clusters (Smith et al., 2018). If this model is accurate, it has an important ramification with respect to the ancestry of the *Hox* segmental patterning functions seen in gnathostomes (Onimaru and Kuraku, 2018; Parker et al., 2016). Since paralogous segmental enhancers exist in gnathostomes, such as the r5 enhancers of *Hoxb3* and *Hoxa3,* and these paralogues are posited to have arisen from duplication before the split between gnathostome and cyclostome lineages, then such segmental regulation presumably also pre-dates this split, as supported by the expression analyses presented here and in previous studies (Parker et al., 2014a; Takio et al., 2007; Takio et al., 2004).

The two additional *Hox* clusters found in lamprey most likely derive from duplication event/s that occurred in the lamprey/cyclostome lineage. In support of this, comparisons of gene content between lamprey Hox-bearing chromosomes suggests that the chromosomes containing the −β and −ε clusters derive from such duplication, as well as those bearing the −α and −δ clusters (Smith et al., 2018). Thus, comparisons between lamprey *Hox* paralogues from the −β and −ε clusters and from the −α and −δ clusters could illuminate patterns of functional divergence that may underlie their retention subsequent to duplication, as discussed below.

### 4.2 The hindbrain hox-code and rhombomeric expression domains

In lamprey, transient rhombomeric segmentation has been described through analyses of morphology, neuro-anatomy and segmental gene expression, being particularly apparent between st22-st24 (Horigome et al., 1999; Kuratani et al., 1998). Segmentally-restricted *Hox* expression, which maps to rhombomere boundaries, had previously been revealed at these stages for three anterior *hox* genes in lamprey: *hoxβ1, hoxα2* and *hoxα3* and compared directly with the expression domains of two genes involved in segmentation (*kreisler* and *krox20)* (Parker et al., 2014a). Collectively, these genes show similar rhombomere-restricted expression domains compared with their gnathostome counterparts, suggesting conservation of a hindbrain gene regulatory network in lamprey. Here, we have expanded upon this initial analysis by demonstrating that all 14 PG1-4 genes are dynamically expressed in the developing hindbrain at the stages examined, with 8 genes exhibiting segmentally-restricted expression at st23: *hoxβ1, −α2, −δ2, −α3, −δ3, −ζ3, −α4* and *−ζ4* (Fig. 8A,C). The six genes lacking segmentally-restricted expression do not have sharp anterior borders and their expression resides in the caudal hindbrain, where segmental markers are not apparent. Electron microscopy and immunolabelling approaches delineated r1-r6 in Arctic lamprey embryos but did not reveal an r6/r7 boundary (Horigome et al., 1999; Kuratani et al., 1998). However, the sharp expression boundaries we identified for *hoxα4* and *hoxζ4* suggest that an r6/r7 boundary exists in lamprey (Fig. 7A), at least at the level of gene expression, and that some of the *Hox* genes are no longer tightly coupled to this segment border (Fig. 8A). From this data, and by comparison with gnathostomes, we can reconstruct aspects of *Hox* expression that were present in the ancestral vertebrate hindbrain and which have been conserved across vertebrates: expression of a PG2 gene up to the r1/r2 boundary, a PG1 gene in r4, elevated expression of a PG3 gene in r5, and expression of a PG4 gene up to the r6/r7 boundary (Fig. 8A,C). Additionally, r1 is devoid of *hox* PG1-4 expression during lamprey hindbrain segmentation; this is also seen in gnathostomes, although *Hox* expression in specific neurons of r1 has been detected at later stages of hindbrain development in some species (McClintock et al., 2003).

**Figure 8:**
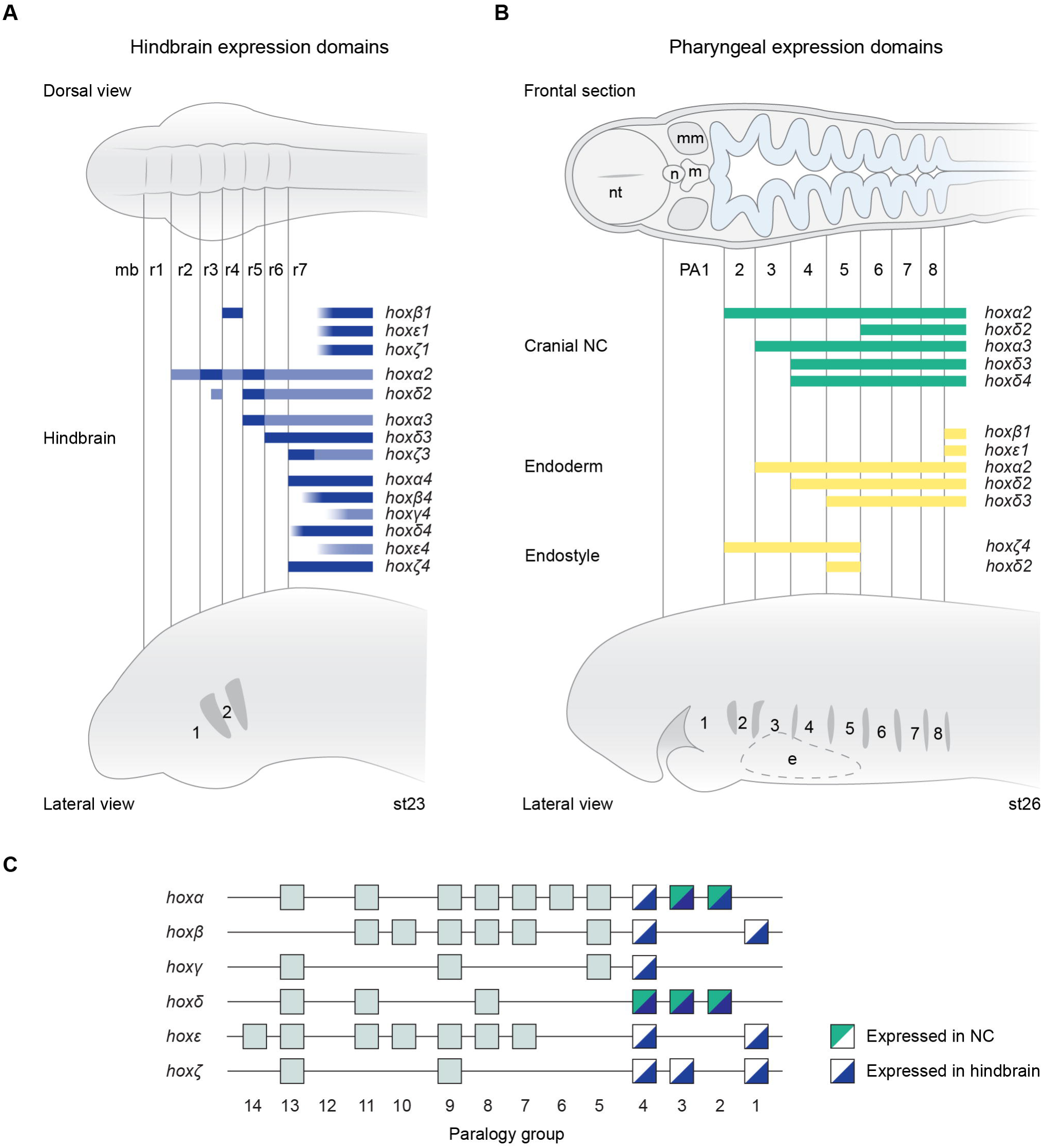
Summary of lamprey *hox*PG1-4 gene expression in the hindbrain and cranial NC. (A) A summary figure depicting segmental domains of expression of lamprey *hox*PG1-4 genes in the hindbrain at st23, shown relative to dorsal (top) and lateral (bottom) schematic representations of the lamprey embryonic head. Rhombomeres (r1-r7) and pharyngeal arches (1-2) are annotated. The blue shading indicates domains of gene expression, with darker shading indicating higher levels of expression as detected by *in-situ* hybridisation. (B) A summary of expression domains of lamprey hoxPG1-4 genes in the cranial NC (green) and endoderm (yellow) at st26, shown relative to schematic representations of the embryonic head in frontal section (top) and lateral view (bottom). Expression in the endoderm-derived endostyle is also shown (yellow). The pharyngeal arches (1-8) are labelled. *hoxβ1* and *hoxε1* have dynamic expression in the endoderm, which retreats caudally during development, being associated with the most recently formed pharyngeal pouch. (C) A depiction of the lamprey *hox* clusters, with *hox*PG1-4 genes marked according to whether they are expressed in NC, hindbrain or both. This reveals that all of the *hox*PG1-4 genes are expressed in the hindbrain, while only genes from *hoxα* and *hoxδ* clusters appear to be expressed in cranial NC at the stages examined, e, endostyle; ec, ectoderm; en, endoderm; m, mouth; mb, midbrain; me, mesenchyme; mm, mandibular mesoderm; n, notochord; nt, neural tube.

A striking aspect of *hox* expression in the arctic lamprey hindbrain is that anterior *hoxα3* domains do not appear to be segmentally restricted at later stages of hindbrain development (st25-st26), despite the maintenance of segmental *krox20* and *ephC* expression (Murakami et al., 2004; Takio et al., 2007). A similar escape from segmental restriction is seen for *hoxα3* in sea lamprey, with rhombomeric registration observed at earlier stages (Fig. 5). Our results reveal that segmental expression perdures through later stages for some lamprey *hox* genes, such as *hoxβ1* and *−ζ4*, while others appear to escape segmental restriction, including *hoxδ2* and *−δ3.* Certain PG4 genes − *hoxγ4, −δ4* and *−ε4* - also exhibit non-uniform anterior expression boundaries at later stages, however it is unclear whether these align with segments, particularly in the caudal hindbrain.

In gnathostome embryos, such escape from segmental registration has not been observed, as once *Hox* expression becomes refined to specific segments and bands of neuronal progenitors over time the domains remain aligned within rhombomere-derived territories during later embryogenesis (Gavalas et al., 2003; Prince et al., 1998; Wingate and Lumsden, 1996). This is regulated in part through a multi-step process whereby early domains are established by c/s-elements that integrate inputs from signaling pathways and the segmental pattern is actively maintained in later stages by auto- and cross-regulatory interactions (Gould et al., 1998; Manzanares et al., 2001; Studer et al., 1998). This suggests that in the lamprey hindbrain there may be key regulatory differences in how and whether *Hox* genes remain coupled to segmentation in later stages, resulting in a temporal relaxation in segmental constraints compared with gnathostomes. This may enable some *hox* genes to be co-opted to perform additional non-segmental roles at later stages of development. Nevertheless, such early segmentation has a lasting effect on the neuronal architecture of the larval lamprey hindbrain, as seen by the segmental arrangement of reticulospinal neurons and the general A-P positioning of cranial nerve motor nuclei (Gilland and Baker, 2005; Murakami et al., 2004; Osorio et al., 2005).

A recent study focusing on hagfish *Hox* genes revealed segmented and nested domains in the embryonic hindbrain, supporting conservation of aspects of this ancestral *Hox* pattern, although differences were also observed, such as the absence of detectable *Hox1* expression from r4 at the stages examined (Pascual-Anaya et al., 2018). Taken together, this points to the existence of an ancient gene regulatory network for Hox-patterning in the hindbrain and pharynx that has been broadly conserved across all vertebrates, but that also exhibits lineage-specific diversification (Parker et al., 2016).

### 4.3 The neural crest Hox-code

The lamprey pharynx comprises 8 pharyngeal arches, which are populated by NC cells migrating in three streams from the hindbrain, broadly equivalent to the three anterior streams of gnathostomes, although a vagal NC stream from the caudal hindbrain appears to be absent in lamprey (Green et al., 2017; Horigome et al., 1999; McCauley and Bronner-Fraser, 2003; Meulemans and Bronner-Fraser, 2002). PA1 is homologous to the mandibular arch of gnathostomes; however, rather than giving rise to the jaw, it forms the velum, a cyclostome-specific piston-like valve involved in ventilating the larval lamprey pharynx (Miyashita, 2016). PA2 is homologous to the hyoid arch of gnathostomes, forming the velar support cartilage, while PA3-8 hold gills, like the posterior pharyngeal arches in aquatic gnathostomes. In gnathostomes, *Hox* PG2-4 genes have nested expression in pharyngeal arch NC that is broadly conserved between species, but paralogues often exhibit differences in expression levels (Parker et al., 2018). Previous studies in lamprey found conservation of select *Hox* domains in NC populations between lamprey and gnathostomes (Takio et al., 2007; Takio et al., 2004). Our data expands on this by showing that none of the PG1-4 genes are expressed in PA1 NC at the stages examined, similar to gnathostomes, and that there are nested domains of expression of five lamprey *hox* genes in PA2-5 (Fig. 8B).

We observe that only genes from *hoxα* and *hoxδ* clusters appear to have nested expression in lamprey cranial NC at the stages examined (Fig. 8C). *hoxα2* is the only PG2 gene expressed in PA2 at these stages, with *hoxα3* the only PG3 gene in PA3, and *hoxδ4* the only PG4 gene in PA4. This suggests that there may be little functional overlap in NC patterning between *hox* genes from the same paralogy group in lamprey. In contrast, some paralogous *Hox* genes share NC expression domains in gnathostome species and exhibit a degree of functional redundancy (e.g. *hoxa2b* and *hoxb2a* in zebrafish PA2, *Hoxa3* and *Hoxb3* in mouse PA3) (Hunter and Prince, 2002; Manley and Capecchi, 1997). This shared activity of paralogues indicates that these *Hox* domains in NC were probably a feature of the ancestral, pre-duplicated *Hox* cluster. If so, then after the *Hox* cluster duplications in ancestral vertebrates, divergent vertebrate lineages have differentially retained NC expression of their paralogous *Hox* genes. However, it is not immediately apparent whether retention versus loss of the NC expression domains of duplicated *Hox* genes has an adaptive significance. Testing the functional roles of lamprey *hox* genes in determining the identity of skeletal elements in the head by CRISPR approaches will be an interesting avenue for future research.

### 4.4 Endodermal hox expression domains

In chick and dogfish, endodermal *Hox* expression has been shown to correlate with specific pharyngeal pouches: *Hoxb1* expression progressively shifts caudally such that it is only present in the most recently formed pharyngeal pouch, *Hoxa2* is associated with the 2^nd^ pharyngeal pouch and *Hoxa3* transiently with the 3^rd^ pouch (Shone et al., 2016). Our analysis reveals dynamic *hoxβ1* and *−ε1* expression in the most recently formed pouch in lamprey (Fig. 3B, 8B), suggesting that the posterior limit of the pharynx is homologous between lamprey and gnathostome species. An RA-dependent role for *Hox1* in defining the posterior limit of the pharynx has been shown in amphioxus (Schubert et al., 2005) and this expression is conserved in a hemichordate, *Saccoglossus kowalevskii* (Gillis et al., 2012), suggesting that this role for *Hox1* genes in pharyngeal development traces its evolution deep into the deuterostome lineage and has been conserved in many extant chordates. Expression was also detected for other *hox* genes in the lamprey pharyngeal endoderm at the stages we examined: *hoxα2* up to the 2^nd^ pouch (st25), *hoxδ2* up to the 3^rd^ pouch (st26) and *hoxδ3* up to the 4^th^ pouch (st26), although this was often at low levels relative to their expression in other domains (Fig. 7, 8B). This suggests that these genes may play similar roles in patterning the pharyngeal endoderm compared to their homologues in gnathostomes.

Among extant vertebrates, lamprey species are unique in possessing an endostyle, which plays a role in filter feeding in larval lampreys and has other functions including regulating iodine uptake. The lamprey endostyle evaginates from endoderm in the ventral pharynx and is transformed into a thyroid during metamorphosis (Kluge et al., 2005). We detected expression in the endostyle for two *hox* genes: *hoxζ4* throughout the A-P extent of the endostyle and *hoxδ2* restricted to the caudal end (Fig. 8B). In urochordates, *Hox1* genes have been implicated in endostyle patterning (Canestro et al., 2008; Yoshida et al., 2017), while *Hox3* genes are required for normal thyroid development in mice (Manley and Capecchi, 1995; Manley and Capecchi, 1998). This suggests that there may be similar requirements for *Hox* genes in patterning these endoderm-derived pharyngeal organs across chordates. However, non-orthologous *Hox* genes appear to be utilised in each of these cases, so further investigation is required to establish whether these reflect conserved ancestral patterning networks or whether this *Hox* patterning has been acquired independently in different lineages.

### 4.5 Patterns of sub-functional divergence between paralogues

Phylogenetic and synteny analyses suggest that the lamprey *hox−β* and *−ε* clusters and the *−α* and *−δ* clusters arose from chromosome-scale duplication event/s in lamprey/cyclostomes, after the gnathostome-cyclostome divergence (Smith et al., 2018). Comparing the PG1-4 gene complement between these duplicated clusters indicates that they have retained *Hox* paralogues to a high degree. This is interesting given the importance of *Hox* genes in development of the body plan and regional specializations. This raises the question of how the lamprey lineage may have utilized these duplicated *hox* genes and the possibilities they may offer in regulating anatomical novelties.

Comparisons of spatiotemporal expression between the pairs of lamprey *hox* paralogues from these duplicated clusters are summarized in Table 1, which illuminate patterns of divergence that may underlie their retention subsequent to duplication. Divergence is seen in the anterior limits of expression between paralogues, such as for *hoxα3* and *hoxδ3* in hindbrain and NC. In other cases, paralogues differ more drastically in tissue specificity, for instance *hoxδ4* retains expression in NC up to PA4, which is presumably ancestral since it is a feature of certain gnathostome PG4 genes, while *hoxα4* has lost expression in this domain. Differences in initiation and maintenance of expression are also seen between paralogues. For example, *hoxβ1* has later onset in the neural plate than *hoxε1* and is maintained in r4 while *hoxε1* expression is lost from this domain (Fig. 3). In mouse, differences in onset and maintenance between *Hoxa1* and *Hoxb1* are attributable to specific c/s-regulatory elements that are associated with each gene: both have a 3’ RARE for early neural expression, while *Hoxb1* is maintained in r4 by an auto-regulatory element that is lacking from *Hoxa1* (Dupé et al., 1997; Marshall et al., 1994; Pöpperl et al., 1995). This suggests that homologous regulatory elements may also have been partitioned between lamprey *hoxβ1* and *hoxε1* after their duplication in the lamprey/cyclostome lineage. Similar patterns of sub-functionalisation or function shuffling with respect to r4 expression have been demonstrated for zebrafish *hoxb1a* and *hoxb1b,* which resulted from the teleost whole genome duplication (McClintock et al., 2001; McClintock et al., 2002). The re-occurrence of this sub-functional partitioning in multiple lineages may be stochastic, or it may reflect features in the organization of *HoxPG1 cis*-regulation such that there is an increased likelihood of sub-functionalisation occurring due to the modularity and functional independence of the enhancer elements that mediate initiation versus maintenance of expression.

**Table 1:** Conserved and divergent expression features of paralogous lamprey *Hox* gene pairs that arose from duplication in the lamprey/cyclostome lineage.

| Paralogous gene pair | Conserved expression features | Divergent expression features |
| --- | --- | --- |
| *hoxβ1, hoxε1* | Pharyngeal endoderm<br>Midbrain neurons<br>Posterior lateral line ganglion | Early onset in neural plate (*hoxε1*)<br>Maintenance in r4 (*hoxβ1*)<br>Anterior lateral line, petrosal and nodose ganglia (*hoxβ1*) |
| *hoxα2, hoxδ2* | PA6-posterior NC | PA2-5 NC (*hoxα2*)<br>Rhombomeric domains<br>Notochord (*hoxδ2*)<br>PA1 mesoderm (*hoxδ2*) |
| *hoxα3, hoxδ3* | r6-posterior<br>PA4-posterior NC | r5 (*hoxα3*)<br>PA3 NC (*hoxα3*)<br>pharyngeal endoderm (*hoxδ3*) |
| *hoxα4, hoxδ4* | Caudal hindbrain<br>Spinal cord | r6/r7 boundary (*hoxα4*)<br>PA4-posterior NC (*hoxδ4*)<br>Maintenance in hindbrain (*hoxδ4*) |
| *hoxβ4, hoxε4* | Caudal hindbrain<br>Spinal cord | Anterior limit in hindbrain |

Did the duplication/s in lamprey/cyclostomes add any new functions to *Hox* genes or simply result in shuffling an ancient set of *Hox* patterning functions such that they became partitioned across duplicated *Hox* clusters? In zebrafish, PG1 and PG5 genes exhibit such partitioning of ancestral functions across paralogues subsequent to the teleost whole genome duplication (Bruce et al., 2001; Jozefowicz et al., 2003; McClintock et al., 2001; McClintock et al., 2003; Prince and Pickett, 2002). In contrast, the lamprey PG2 genes may represent a different case, with more dramatic differences in expression between paralogues. These include expression domains for *hoxδ2* that, to our knowledge, have not been seen for PG2 genes in other vertebrate species, such as in PA1 mesoderm and in the endostyle (Table1). These may reflect ancestral vertebrate *Hox2* functions that have been retained in lamprey but lost in gnathostomes. However, such expression domains have not been characterized for invertebrate deuterostome *Hox2* genes, so it is unclear whether they are ancestral to vertebrates. Another intriguing possibility is that these expression domains evolved in the lamprey/cyclostome lineage, after duplication of the *hoxα* and *−δ* clusters (neo-functionalisation). Coupled with these non-canonical expression domains, *hoxδ2* genes in both sea lamprey and arctic lamprey show a high degree of sequence divergence relative to other vertebrate *Hox2* genes (Pascual-Anaya et al., 2018). This may reflect either relaxation of selective constraint or positive selection for discrete functions after duplication in the lamprey/cyclostome lineage. It will be interesting to address the functional significance of these expression domains.

### 4.6 Conclusion

In conclusion, our analysis suggests that many *Hox* expression domains that are observed in extant gnathostomes were present in ancestral vertebrates but have been partitioned differently across *Hox* clusters in gnathostome and cyclostome lineages after duplication. On top of this conserved *Hox* patterning ground-plan, lamprey also shows differences in spatiotemporal *Hox* expression, which may or may not be ancestral. These include tissue domains that are either not present or not associated with *Hox* expression in gnathostomes, such as the endostyle and PA1 mesoderm. Understanding how these conserved and divergent *Hox* expression domains relate to vertebrate head evolution will require examination of *Hox* functional roles in lamprey using CRISPR knockout approaches (Square et al., 2015). Such approaches could test the assumption that segmental *Hox* expression plays equivalent roles in lamprey and gnathostomes and could address the functional significance of the differences in *Hox* expression that we observe. Characterization of lamprey *Hox* enhancer elements and comparison with those of gnathostomes will enable inference of common ancestral *Hox* regulatory mechanisms in vertebrates, and may elucidate how *Hox* functions have been differentially partitioned across *Hox* clusters in lamprey versus gnathostome lineages (Parker et al., 2014b). Looking deeper in chordate evolution, these studies will provide a platform for regulatory comparisons with non-vertebrate deuterostomes (Minor et al., 2018), to investigate how vertebrate segmental *Hox* regulation arose.

## Acknowledgements

We thank Stephen Green, Dorit Hockman, Tetsuto Miyashita, and Megan Martik for lamprey husbandry assistance, and the Stowers Institute Histology facility for sectioning assistance. HJP and RK were supported by the Stowers Institute (RK grant #2013-1001). MEB was supported by grants R01NS108500 and R35 NS111564.

